# Multi-center Translational Trial of Remote Ischemic Conditioning in Acute Ischemic Stroke (TRICS BASIC)

**DOI:** 10.1101/2025.03.25.645373

**Authors:** Simone Beretta, Mauro Tettamanti, Jacopo Mariani, Susanna Diamanti, Alessia Valente, Ornella Cuomo, Chiara Di Santo, Ilaria Dettori, Martina Venturini, Irene Bulli, Elisabetta Coppi, Manuel Alejandro Montano Castillo, Erica Butti, Giorgia Serena Gullotta, Martina Viganò, Francesco Santangelo, Francesco Andrea Pedrazzini, Carlo Perego, Serena Seminara, Serenella Anzilotti, Joanna Rzemieniec, Laura Castiglioni, Benedetta Mercuriali, Majeda Muluhie, Chiara Paola Zoia, Gessica Sala, Luigi Sironi, Marco Bacigaluppi, Gianvito Martino, Felicita Pedata, Anna Maria Pugliese, Diana Amantea, Giacinto Bagetta, Antonio Vinciguerra, Giuseppe Pignataro, Stefano Fumagalli, Maria-Grazia De Simoni, Carlo Ferrarese

## Abstract

**Background:** Basic science studies have reported remote ischemic conditioning (RIC) as neuroprotective in acute ischemic stroke, while clinical evidence remains conflicting. The TRICS BASIC study investigated the efficacy and safety of RIC in experimental ischemic stroke using a rigorous clinical trial methodology.

**Methods:** Multi-center, multi-species, parallel group, randomized, controlled, preclinical trial of transient femoral artery clipping to induce RIC in female and male rats and mice subjected to transient endovascular occlusion of the middle cerebral artery. Animals were randomized to receive RIC, or sham surgery, after reperfusion. The primary endpoint was good functional outcome at 48 hours, assessed using a composite functional neuroscore. Secondary endpoints was infarct volume at 48 hours and safety, assessed using a standardized health report at 24 and 48 hours. Pre-enrollment harmonization, centralized monitoring, allocation concealment, blinded outcome assessment and intention-to-treat analysis were applied.

**Results:** The trial enrolled 164 rodents (82 mice and 82 rats) of both sexes (53% females), across seven laboratories. A greater proportion of RIC-treated rodents achieved a favorable functional outcome compared to controls, at 48 hours post-ischemia (55% versus 36%; OR 2.2, 95% CI [1.23–4.4], p=0.009). RIC was associated with a small reduction in infarct volume (standardized mean difference -0.38, 95% CI [−0.70, −0.05], p=0.024). Health monitoring indicated no major safety concerns, and post-operative analgesia requirements were lower in RIC-treated mice.

**Conclusions:** Surgically-induced RIC provided a modest but evident neuroprotective effect in experimental ischemic stroke, underscoring the potential of this strategy as an adjunctive treatment in stroke care. The findings of the TRICS BASIC study highlighted the importance of multicenter preclinical trials in addressing variability and enhancing translational validity.

**Registration:** registered at preclinicaltrials.eu, identifier PCTE0000177.

## INTRODUCTION

Stroke is one of the leading causes of death and disability at a global level ^1^, with the only effective treatment being time-dependent recanalization of cerebral arteries through intravenous thrombolysis or endovascular mechanical thrombectomy ^2^.

The economic impact of cardiovascular risk factors is anticipated to rise substantially in the coming decades, concomitant with a projected increase in the incidence of stroke and stroke- related costs ^3^. This underscores an urgent need to develop and integrate novel therapeutic strategies. Such approaches could complement current recanalization therapies or serve as alternatives in cases where these interventions are contraindicated, with the ultimate goal of improving clinical outcomes in stroke management.

The development of neuroprotective strategies has encountered significant translational barriers, with the majority of candidate therapies demonstrating promising preclinical efficacy but failing to achieve successful outcomes in clinical trials ^4^. To bridge this gap, the concept of multi-center preclinical randomized controlled trials (RCTs) has emerged as a critical intermediary step, mirroring the rigor and design of phase 3 clinical trials and addressing variability across experimental settings ^5,6^.

Remote ischemic conditioning (RIC) is a promising neuroprotective strategy that aims to induce systemic ischemic tolerance through brief episodes of ischemia and reperfusion applied to distant organs. Initially demonstrated in the heart, RIC has been shown to activate endogenous protective mechanisms against ischemic brain injury ^7–10^.

The encouraging results observed in single-laboratory studies ^11,12^, require confirmation through multi-center preclinical RCTs to robustly evaluate the efficacy and safety of RIC in stroke models.

Currently completed phase 2 and 3 clinical RCTs on RIC in patients with acute ischemic stroke were conducted using different application protocols and provided conflicting results ^13–20^.

The TRICS BASIC multi-center preclinical RCT was designed to investigate efficacy and safety of surgical RIC, by transient femoral artery clipping, in two animal species (mice and rats) and both sexes, using the stroke model of transient middle cerebral artery (MCA) occlusion. This preclinical trial was realized within a network of seven Italian laboratories with extensive expertise in the transient MCA occlusion model, aiming to generate high-quality, reproducible data to inform future clinical translation.

## METHODS

### Study design

The study protocol was pre-registered on preclinicaltrials.eu (Identifier: PCTE0000177). Detailed methodology of the TRICS BASIC study has been published as a protocol paper ^21^ before starting randomization of experimental animals.

The data that support the findings of this study are available from the corresponding author upon reasonable request.

Authorization for animal use was granted under project licenses issued by the Italian Ministry of Health (1056/2020-PR). All experimental procedures adhered to national regulations governing the use of laboratory animals (D.L. 26/2014) and conformed to the European Union Directive on the protection of animals used for scientific purposes (2010/63/EU). The experimental design and conduct were aligned with the National Institutes of Health guidelines and complied with the Animal Research: Reporting of In Vivo Experiments (ARRIVE) guidelines to ensure transparency and reproducibility in animal research.

The TRICS BASIC study was designed as a multicenter, multispecies, randomized, controlled, preclinical trial, to investigate efficacy and safety of RIC in experimental ischemic stroke and was carried out in seven Italian centers between September 2021 and January 2023.

There were two successive randomizations: (1) allocation to receive an occlusion of the middle cerebral artery (MCA+) or its sham equivalent (MCA-), with a 5:1 ratio, and (2) allocation to receive a remote ischemic conditioning (RIC+) via transient femoral artery occlusion, or its sham equivalent (RIC-), with a 1:1 ratio.

To ensure allocation concealment, each center received sealed, non-transparent, and non- resealable envelopes for the 2-steps randomization, marked with codes for species (mouse or rat), sex, and progressive numbers. Upon opening an envelope, a sheet detailing center identification, sex, and MCA+/MCA- assignment was signed and dated. A second internal envelope containing the RIC+/RIC- treatment was opened only after MCA surgery to ensure allocation concealment for surgeons. Randomization lists and envelopes were prepared by personnel not involved in animal procedures and securely stored.

Animals of both sexes were equally represented. Animals which either scored below the prespecified threshold of 2 of the intraischemic clinical score (see below) or with no ischemic lesion detected at brain histology were excluded from the analyses (i.e. animals without stroke). Animals that died before RIC randomization were excluded (i.e.: animals not reaching the target randomization) and replaced by other animals, up to three per center. Conversely, animals that died after RIC randomization were retained in the prespecified (intention-to-treat) primary analyses and given the worst score: this was true for naturally dying animals and for animals sacrificed after showing signs of extreme distress.

### Middle cerebral artery occlusion in mice and rats

Adult C57BL/6J mice (weight 26 g ± 5%) and adult Sprague-Dawley rats (weight 250 g±5%), male and female 1:1 (regardless of oestrous cycle), housed in single cages, exposed to 12/12 hours light/dark cycle, at controlled room temperature, with free access to food and water, in specific pathogen-free facilities were used. Animals were anesthetized by 3% isoflurane in O2/N2O (1:3) and maintained by 1%–1.5% isoflurane in O2/N2O (1:3). Transient occlusion of the origin of the right MCA was induced (60 min in mice, 100 min in rats) followed by a reperfusion period of 48 hours. In mice, a silicone-coated monofilament nylon suture (sized 7–0, diameter 0.06–0.09 mm, length 10±1 mm; diameter with coating 0.23 mm; coating length 2 mm, Doccol Corporation, Redlands, California, USA) was introduced into the right common carotid artery and advanced to block the right MCA. In rats, a silicone-coated filament (sized 5–0, diameter with coating 0.33 mm; coating length 5–6 mm; Doccol Corporation, Redlands, California, USA), was introduced in the right external carotid artery and pushed through the internal carotid artery to occlude the origin of the right MCA.

For both species, body temperature was kept at 37°C by a heating pad during surgery. During MCA occlusion, animals were awakened from anaesthesia, kept in a warm box and tested for the intraischemic clinical score (see below) in single cages. After 60 min, blood flow was restored by carefully removing the filament, under anesthesia. Sham rodents received the same anesthetic regimen and surgery than MCA occluded animals, i.e. their common, external and internal carotid arteries were isolated, but the filament was not introduced. After surgery, animals were returned to single cages.

### Intraischemic clinical score

After MCA occlusion, the following intraischemic clinical score was applied to test the correct induction of cerebral ischemia. Animals were judged ischemic and included in the trial, if presenting 3 or more of the following deficits after filament insertion:

1. The palpebral fissure had an ellipsoidal shape (not the normal circular one).
2. One or both ears extended laterally.
3. Asymmetric body bending on the ischemic side.
4. Limbs extended laterally and did not align to the body.

### Remote ischemic conditioning in mice and rats

RIC was induced via transient femoral artery occlusion. At a set time after reperfusion (20 minutes in rats, 10 minutes in mice), the femoral artery was identified, isolated, and transiently occluded with a microsurgical clip to halt blood flow for the specified duration (20 minutes in rats, 10 minutes in mice). Successful occlusion was confirmed by visually inspecting the distal femoral artery. Sham-treated animals (RIC-) underwent the same procedure without femoral artery clipping.

### Outcomes

Outcomes were measured at the end of the study (48 hours after MCA occlusion).

The primary outcome was the proportion of animals (separately by species) with good functional outcome assessed by the De Simoni neuroscore ^22,23^, which comprehensively assessed global and focal sensorimotor deficits. Good outcome was defined as a De Simoni neuroscore of 20 or less, on a scale from 0 (no deficits) to 56 (all deficits present at maximum value).

The coordinating unit conducted centralized training on the correct administration of the De Simoni composite neuroscore through multiple pre-randomization meetings and a detailed instructional video ^24^.

Secondary outcomes were:

- infarct volume measured by volumetric histology; brains were fixed in ice-cold 10% neutral buffer formalin and shipped to the coordinating center (University of Milano-Bicocca) for centralized processing and blinded evaluation; coronal sections (50 µm) were obtained and stained using Cresyl Violet 0.1% (Bioptica, Milano, Italy). Infarct areas were measured in consecutive sections with 250 µm interval (on average, 19 sections for rat brain and 9 sections for mouse brains). Infarct volume was calculated using ImageJ image processing software (National Institute of Health, Bethesda, MD, USA), corrected for inter-hemispheric asymmetries due to cerebral edema, and expressed in mm^3^.
- De Simoni neuroscore, considered as a continuous value.

### Blinding

Functional and clinical assessments were conducted locally by researchers blinded to MCA occlusion and RIC treatment allocation, i.e. by a person not participating in/assisting the surgical practices. Centrally conducted histological procedures and data analysis (RIC+ vs RIC-) were conducted blinded to the group allocation.

### Data collection

Study data were collected and managed using REDCap electronic data capture tools hosted at the Istituto di Ricerche Farmacologiche Mario Negri IRCCS on behalf of the coordinating unit.

REDCap served as a secure web-based platform for research data management, offering validated data entry, audit trails, automated data exports, and integration with external sources for statistical analysis.

### Health and safety assessment

An “MCAO health and safety report” documented data on animal identification, procedural details, and post-surgical outcomes. Evaluations occurred pre-surgery, intraoperatively, and at 24 and 48 hours post-surgery, following the IMPROVE guidelines ^25^. Key metrics included weight changes, isoflurane exposure, respiratory rates, surgical times, and postoperative distress categorized as mild, moderate, or severe. This systematic approach ensured comprehensive tracking of health parameters and adverse outcomes to promote animal welfare during the conduct of the study.

### Sample size calculation

A 20% rate of good functional outcomes was used as the baseline for animals subjected to MCA occlusion without effective treatment. An improvement of at least 30% (from 20% to 50%) was established as the minimum effect size of translational significance. Statistical significance (α) was set at 0.050 (two-tailed). Using a chi-squared test for data analysis, a total of 80 animals equally randomized between RIC+ and RIC- groups yielded a statistical power of 82%. The same calculation was applied to both species, requiring 160 animals to undergo MCA occlusion. Accounting for a 30% exclusion rate, a target of 120 animals randomized per species was set. No power calculation was conducted for non-occluded animals, as they served solely as internal controls.

### Statistical methods

Descriptive analyses were conducted on baseline and procedural characteristics. The primary outcome, the dichotomized De Simoni neuroscore, was analyzed using a logistic regression model, which included randomization strata (RIC, center [as a random effect], and sex). To explore potential sex-specific effects of RIC, models incorporating a sex-by-RIC interaction were also tested, but were not selected as the final models due to non-significant results (p>0.20 from likelihood ratio tests). An intention-to-treat approach was applied, with animals that died after RIC being assigned a negative outcome. The two parallel trials were analyzed separately by species, and results were combined in an overall analysis if one trial showed a borderline significant p-value.

Secondary outcomes (infarct volume and continuous De Simoni neuroscore) were analyzed using mixed linear regressions, with sex and RIC as covariates, and center as a random effect. To address species differences in infarct size, the infarct volume was expressed as a percentage of the ipsilateral hemispheric volume and standardized by subtracting the group mean and dividing by the group standard deviation.

Health and safety monitoring data were presented as numbers and percentages for the two groups.

Comparisons were made in the MCA+ groups, while the MCA- (sham) group was analyzed descriptively. All analyses were conducted blind using Stata/IC v. 15 (Statacorp).

## RESULTS

### Study flow diagram

Between October 2021 and January 2023, a total of 216 animals (110 mice and 106 rats) were utilized. Of these, 52 animals were excluded based on eligibility criteria, which included death prior to randomization, major surgical complications, randomization to sham surgery, or absence of visible ischemic injury on histological analysis. This resulted in a final sample of 164 randomized animals (82 mice and 82 rats). The study flow diagram is shown in Figure 1.

**Figure 1.**
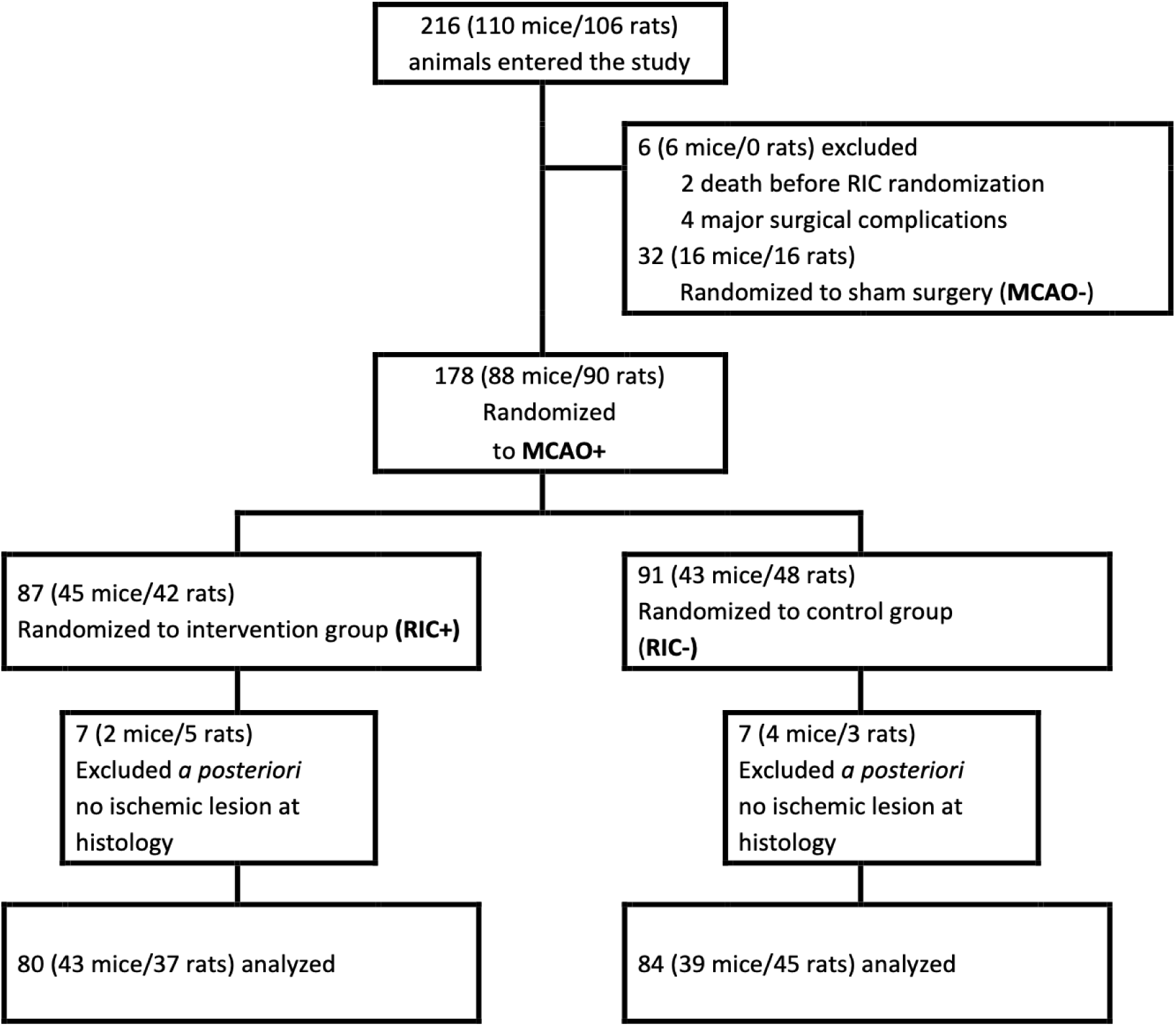
Flow diagram of the TRICS BASIC preclinical trial. MCAO- = sham carotid artery surgery. MCAO+ = middle cerebral artery occlusion. RIC+ = remote ischemic conditioning. RIC- = sham femoral artery surgery.

### Baseline characteristics of the study population

The baseline characteristics of the 164 animals included were well-balanced across treatment groups in terms of sex, weight, body temperature during MCAO or RIC surgery, isoflurane exposure duration, and respiratory rate throughout the surgical procedure (Table 1). Intra- ischemic clinical assessments demonstrated a 92% accuracy in identifying ongoing cerebral ischemia, correctly diagnosing ischemia in 164 animals, while yielding a false-positive rate of 8% (14 animals with no ischemic lesions confirmed by histology; Table 2). As pre-specified in the study protocol, animals with false-positive diagnosis of ischemia were excluded from the final analysis.

**Table 1.** Baseline characteristics of the study population. RIC = remote ischemic conditioning. IQR = interquartile range. n = number.

|  |  | **Mice** |  |  | **Rats** |  |  |
| --- | --- | --- | --- | --- | --- | --- | --- |
|  | unit | Intervention<br>group (RIC+)<br>median (IQR)<br>or n(%) | Control<br>group (RIC-)<br>median (IQR)<br>or n(%) | p value | Intervention<br>group (RIC+)<br>median (IQR)<br>or n(%) | Control<br>group (RIC-)<br>median (IQR)<br>or n(%) | p value |
| **Number** | n | 43 | 39 |  | 37 | 45 |  |
| **Sex**<br>(males/females) | n (%) | 19/24 (44/56) | 18/21 (46/54) | 0.86 | 18/19 (49/51) | 22/23 (49/51) | 0.98 |
| **Weight** |  |  |  |  |  |  |  |
| 24 hours before<br>surgery | gr | 23 (22.2; 24) | 22.9 (22.1; 24) | 0.90 | 247 (230; 271) | 260 (230; 274) | 0.52 |
| at the time of<br>surgery | gr | 23.1 (22.2; 24) | 23 (22.4; 24) | 0.98 | 250 (230; 276) | 262 (230; 285) | 0.65 |
| **Body temperature** |  |  |  |  |  |  |  |
| during MCAO<br>surgery (min) | °C | 36 (35.9; 36.4) | 36.2 (35.9; 36.6) | 0.40 | 36.8 (36.4;<br>37.2) | 36.7 (36.3; 36.9) | 0.53 |
| during MCAO<br>surgery (max) | °C | 37.1 (36.7;<br>37.4) | 37.2 (36.8; 37.6) | 0.31 | 37.5 (37.1;<br>37.7) | 37.2 (37.1; 37.4) | 0.03 |
| during RIC surgery<br>(min) | °C | 36.2 (35.6;<br>36.4) | 36.1 (35.4; 36.5) | 0.73 | 36.9 (36.2;<br>37.3) | 37 (36.1; 37.1) | 0.63 |
| during RIC surgery<br>(max) | °C | 36.9 (36.4;<br>37.4) | 37.2 (36.7; 37.4) | 0.14 | 37.5 (37.3;<br>37.9) | 37.3 (37; 37.8) | 0.22 |
| **Isoflurane**<br>**exposure** |  |  |  |  |  |  |  |
| during MCAO<br>surgery | min | 18 (17; 23) | 18 (13; 23) | 0.95 | 34 (10; 75) | 42 (28; 81) | 0.53 |
| during RIC surgery | min | 27 (24; 32) | 26 (21; 31) | 0.43 | 48.5 (30; 54.5) | 29 (27; 55) | 0.05 |
| **Breath rate** |  |  |  |  |  |  |  |
| at baseline | breaths/<br>minutes | 212 (188; 254) | 204 (76; 248) | 0.55 | 80 (68; 100) | 81 (69; 100) | 0.89 |
| 15 minutes after<br>starting anesthesia | breaths/<br>minutes | 108 (78; 120) | 114 (72; 134) | 0.71 | 60 (56; 72) | 60 (52; 72) | 0.80 |
| 15 minutes after<br>withdrawing<br>anesthesia | breaths/<br>minutes | 158 (106; 188) | 162 (74; 182) | 0.97 | 76 (68; 88) | 74 (64; 88) | 0.70 |

**Table 2.** Performance of the intraischemic clinical score. MCAO- = animals either with sham surgery or subjected to middle cerebral artery occlusion without ischemic lesion at histology. MCAO+ = animals subjected to middle cerebral artery occlusion and ischemic lesion confirmed at brain histology.

|  | **MCAO-**<br>**No ischemic lesion at histology** | **MCAO+**<br>**Ischemic lesion confirmed at histology** |
| --- | --- | --- |
| **Negative intraischemic score (0-2)** | 32<br>True negative 100% | 0<br>False negative 0% |
| **Positive intraischemic score (3-4)** | 14<br>False positive 8% | 164<br>True positive 92% |

### Primary efficacy outcome

Primary outcome data are shown in Table 3 and Figure 2A. The percentage of mice achieving a favorable functional outcome at 48 hours, defined as a score of 20 or less in the De Simoni composite neuroscore, was higher in the RIC+ group (49%) compared to the RIC− group (28%), but this difference reached a borderline significance level in the unadjusted analysis (absolute difference 21%, OR 2.5, 95% CI [0.97–6.1], p=0.058) or when stratified by sex (p=0.052).

**Figure 2.**
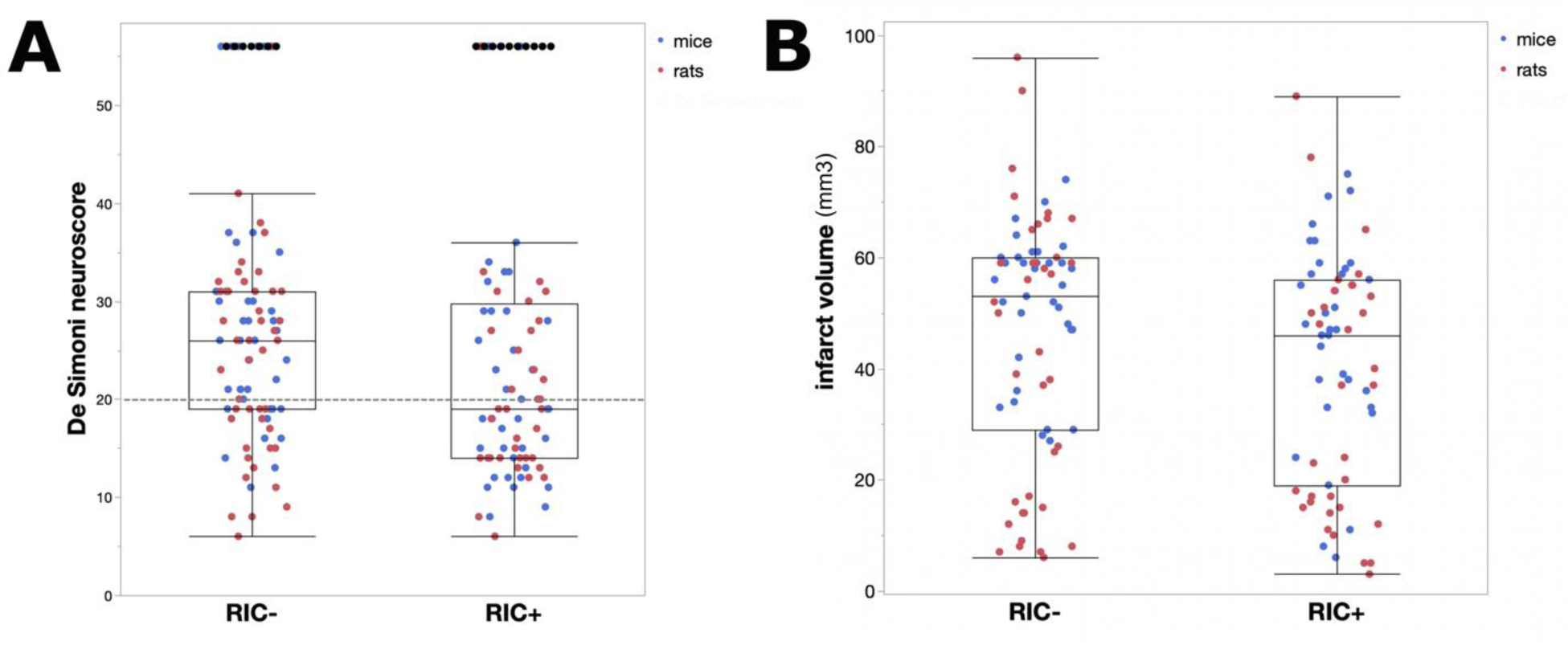
Functional neuroscore and infarct volume. Distribution of De Simoni neuroscore (A) and infarct volume (B) of individual animals enrolled in the TRICS BASIC trial (mice in blue; rats in red). The threshold of 20 for “good functional outcome” is indicated with a dotted line. Data are expressed as box and whiskers plots (median, 25%-75% quartiles, minimum [25% quartile – 1.5*interquartile range], maximum [75% quartile + 1.5*interquartile range], with outliers).

**Table 3.**
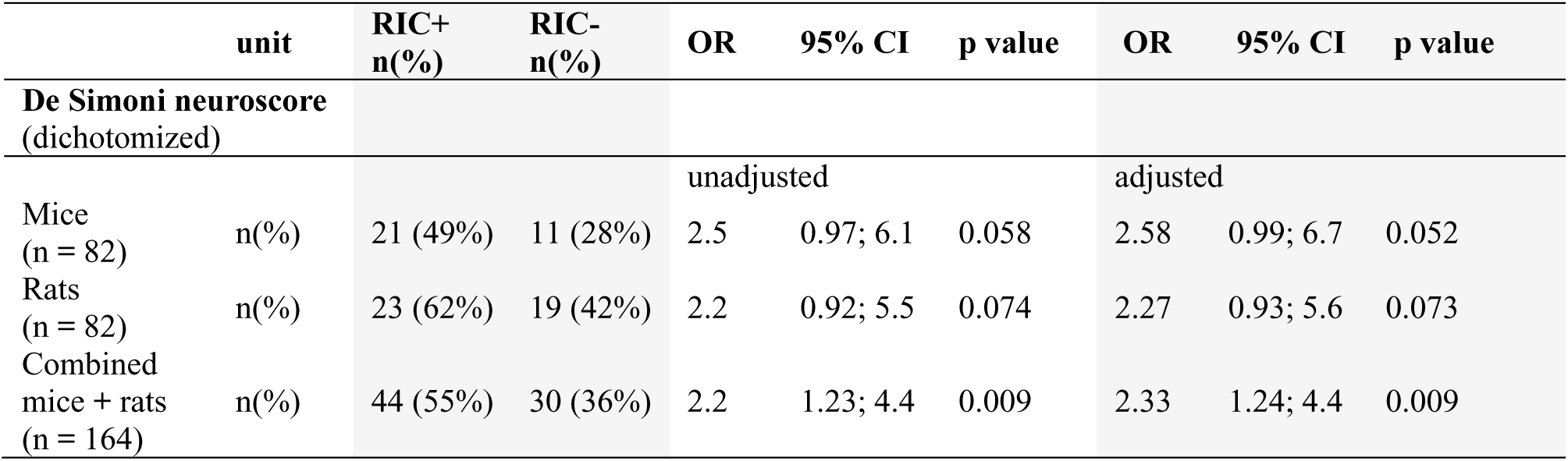
Primary outcome: good (20 or less) versus poor (> 20) De Simoni neuroscore at 48 hours. RIC+= animals treated with remote ischemic conditioning. RIC-= animals with sham femoral artery surgery. OR = odds ratio. CI = confidence intervals. n = number.

Similarly, the proportion of rats with a favorable functional outcome at 48 hours was higher in the RIC+ group (62%) compared to the RIC− group (42%), but this difference also did not reach statistical significance in the unadjusted analysis (absolute difference 20%, OR 2.2, 95% CI [0.92–5.5], p=0.074) or in the sex-stratified analysis (p=0.073). When combining data from both species, a statistically significant benefit was observed for the RIC+ group (55%) compared to the RIC− group (36%), both in the unadjusted analysis (absolute difference 19%, OR 2.2, 95% CI [1.23–4.4], p=0.009) and when including sex as a covariate (p=0.009).

Results of the primary outcome of animals randomized to sham surgery are shown in Figure S1. Results of the primary outcome by single laboratories is shown in Figure S2.

### Secondary Efficacy Outcomes

When the De Simoni neuroscore was considered as a continuous variable (Table 4), mice in the RIC+ group had significantly lower mean values compared to the RIC− group in both the overall analysis (20.1 vs. 24.3; mean difference −4.2, 95% CI [−7.8, −0.50], p=0.027) and the sex-stratified analysis (p=0.022). Similarly, rats in the RIC+ group exhibited significantly lower mean De Simoni neuroscore values compared to the RIC− group in both the overall analysis (19.1 vs. 24.3; mean difference −4.3, 95% CI [−8.0, −0.57], p=0.024) and the sex-stratified analysis (p=0.018). Combining results across species confirmed the benefit of RIC in both the non-sex-stratified analysis (mean De Simoni neuroscore: 19.6 in RIC+ vs. 23.8 in RIC−; mean difference −4.2, 95% CI [−6.8, −1.6], p=0.002) and when sex was included as a covariate (p=0.001).

**Table 4.**
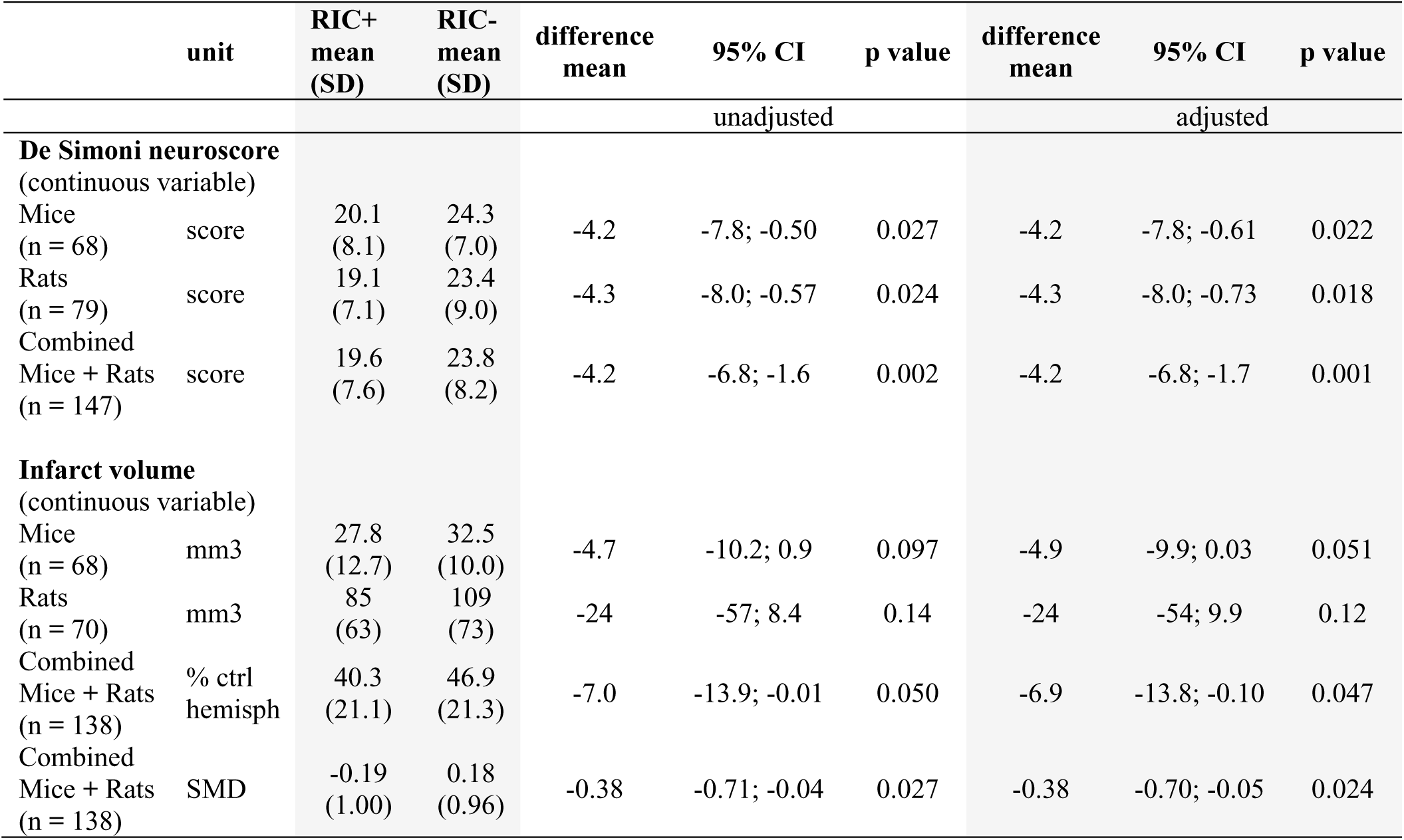
Secondary outcomes: volumetric histology and De Simoni neuroscore as a continuous variable. RIC+= animals treated with remote ischemic conditioning. RIC-= animals with sham femoral artery surgery. SD = standard deviation. CI = confidence intervals. n = number. SMD = standardized mean difference.

The mean infarct volume at 48 hours (Table 4 and Figure 2B) did not significantly differ in RIC+ mice compared to RIC− mice in the unadjusted analysis (27.8 vs. 32.5 mm³; mean difference −4.7, 95% CI [−10.2, 0.9], p=0.097), but this difference reached borderline significance in the sex-stratified analysis (p=0.051). The mean infarct volume at 48 hours was lower in RIC+ rats compared to RIC− rats in the overall analysis, though not statistically significant (85.0 vs. 109 mm³; mean difference −24, 95% CI [−57, 8.4], p=0.14), with similar results in the sex-stratified analysis (p=0.12). The combined analysis of both species, considering the percentage of ischemic brain tissue relative to the total ipsilateral hemisphere, showed that infarct volume was lower in RIC+ animals compared to RIC− animals with borderline significance in the overall analysis (40.3% vs. 46.9%; mean difference −7.0, 95% CI [−13.9, −0.01], p=0.050) and in the sex-stratified analysis (p=0.047). Adjusted standardized mean difference (SMD) was also calculated for infarct volume and was consistent with the volumetric data (-0.38, 95% CI [−0.70, −0.05], p=0.024).

### Safety and Health Monitoring

The health and safety assessments are summarized in Table 5. At 48 hours post-surgery, weight loss ≤10% was observed in 22 mice (27%) and 38 rats (46%), with no significant differences between treatment groups in either species. Weight loss >10% at 48 hours was documented in 47 mice (57%) and 41 rats (50%), similarly showing no treatment group differences.

**Table 5.** Health and safety report of the study population. RIC+= animals treated with remote ischemic conditioning. RIC-= animals with sham femoral artery surgery. IQR = interquartile range. n = number. SMD = standardized mean difference.

|  | unit | Mice |  |  | Rats |  |  |
| --- | --- | --- | --- | --- | --- | --- | --- |
|  |  | Intervention group (RIC+) median (IQR) or n(%) | Control group (RIC-) median (IQR) or n(%) | p value | Intervention group (RIC+) median (IQR) or n(%) | Control group (RIC-) median (IQR) or n(%) | p value |
| **Number** | n | 43 | 39 |  | 37 | 45 |  |
| **Death** |  |  |  | 0.66 |  |  | 0.45 |
| < 24 hours from surgery | n (%) | 7 (16%) | 5 (13%) |  | 2 (5%) | 1 (2%) |  |
| < 48 hours from surgery | n (%) | 8 (19%) | 6 (15%) |  | 2 (5%) | 1 (2%) |  |
| **Weight loss** |  |  |  | 0.60 |  |  | 0.49 |
| 10% or less at 48 hours | n (%) | 11 (31%) | 11 (32%) |  | 19 (54%) | 19 (43%) |  |
| > 10% at 48 hours | n (%) | 24 (56%) | 23 (59%) |  | 16 (43%) | 25 (55%) |  |
| **Distress within 48 hours** |  |  |  | 0.13 |  |  | 0.57 |
| No distress | n(%) | 0 | 0 |  | 1 (3%) | 0 |  |
| Low distress | n (%) | 5 (12%) | 0 |  | 17 (46%) | 19 (42%) |  |
| Moderate distress | n (%) | 7 (16%) | 10 (26%) |  | 13 (35%) | 6 (36%) |  |
| Severe distress | n (%) | 23 (53%) | 23 (59%) |  | 4 (11%) | 9 (20%) |  |
| **Analgesic drug within 48 hours** | n (%) | 13 (30%) | 24 (62%) | 0.004 | 0 | 0 | - |
| **Euthanasia within 48 hours** | n (%) | 1 (2%) | 0 | 0.34 | 0 | 0 | - |

No statistically significant differences were identified in distress severity at 48 hours, ranging from no distress to death, between RIC+ and RIC− groups in both mice and rats. Post-operative analgesia use was significantly higher in the control (RIC−) group compared to the treatment (RIC+) group in mice (62% vs. 30%; *p* = 0.004). Notably, no post-operative analgesia was required for rats in either treatment group. Euthanasia was performed in only one mouse within the RIC+ group.

Mortality rates were comparable across treatment groups. Early mortality (<24 hours post- surgery) occurred in 12 mice (15%) and 3 rats (4%). An additional mouse in each treatment group died during the 24 to 48-hour period, prior to study completion.

## DISCUSSION

The TRICS BASIC study demonstrated that RIC exerts a modest yet potentially significant neuroprotective effect in the experimental stroke model of MCA occlusion and reperfusion. This finding was robustly confirmed in a large multicenter preclinical trial, employing two rodent species of both sexes, and conducted across seven independent research laboratories.

The trial was designed to implement the features required for a clinical trial, including pre- registration, publication of a protocol paper ^21^, sample size estimate, randomization, allocation concealment, centralized monitoring, investigators’ training and blinded outcome assessment. Rodents entering the study were registered into an electronic case report form, where detailed physiological and experimental parameters were recorded before, during and after surgery. The overall methodology of the TRICS BASIC study was conceived with the aim of setting a benchmark for preclinical rigor, similarly to the Stroke Preclinical Assessment Network (SPAN) initiative ^26^.

The inclusion of 216 animals with stringent inclusion/exclusion criteria ensured the reliability of the findings, with a high success rate (92%) in inducing ischemia verified through histology.

Our results robustly indicate that RIC improved functional outcomes and reduced infarct volume after MCA occlusion and reperfusion, albeit to a smaller extent than previously reported.

Specifically, rodents receiving RIC were more likely to achieve a good outcome, defined as a De Simoni neuroscore <20 ^21^. This dichotomous primary outcome was conceptually aligned with the modified Rankin scale 0-2, which is commonly used as the primary outcome measure in clinical stroke trials ^27^. Functional outcome, instead of infarct volume, was purposely chosen as the primary outcome in order to enhance the translational potential of this preclinical trial.

Notably, the De Simoni neuroscore at 48 hours after stroke was reported to correlate with infarct size ^22^. Accordingly, the observed improvement in sensorimotor deficits was associated with a small reduction in the infarct volume. This finding is consistent with the hypothesis that RIC exerts a neuroprotective effect, mitigating both behavioral impairments and structural brain damage.

However, the treatment effect size of RIC on infarct volume was considerably smaller in the TRICS BASIC trial compared to a large preclinical data meta-analysis which reported infarct size as the primary outcome measure (SMD -0.38 versus -2.0, respectively) ^11^.

Secondary analyses bolster these findings. When analyzed as a continuous variable, neuroscores revealed a significant mean improvement in RIC-treated rodents, further supported by a reduction in post-operative analgesic use. These data reinforce the translational relevance of the neuroscore cut-off at 20 and highlight the role of RIC in enhancing overall animal welfare. The consistency of physiological parameters across experimental groups rules out confounding effects, adding confidence to the validity of the observed outcomes.

Notably, the study also addressed variability across centers. While some variability was observed in RIC efficacy, untreated ischemic animals (RIC-) showed consistent neuroscores across all centers, underscoring the robustness of the MCA occlusion model. Variability in RIC outcomes, such as reduced efficacy in certain centers, may reflect differences in technical expertise or baseline methodological variations. Such variability emphasizes the importance of multicenter trials in capturing real-world complexities, thereby enhancing the replicability and generalizability of findings. Importantly, a multi-step, online harmonization phase was conducted before starting enrollment of randomized animals in the TRICS BASIC trial, to implement the De Simoni neuroscore across all participating laboratories, and mitigated the variability in behavioral evaluations before the interventional phase ^24^.

The TRICS BASIC study employed two rodent species and included both sexes, following the recommendations of the RIGOR guidelines for confirmatory preclinical studies ^28^, to enhance the generalizability of results and the translational potential for clinical application.

Clinical evidence regarding the efficacy of RIC in acute ischemic stroke remains inconclusive, potentially due to variations in application protocols. Notably, the RICAMIS trial ^15^ demonstrated a clinical benefit of RIC, whereas the RESIST ^14^, RESCUE BRAIN ^20^, REPOST ^16^, and RECAST-2 ^18^ trials reported neutral results. Of interest, the RICAMIS trial employed a higher dose intensity and extended duration of RIC treatment (5 cycles of 5 minutes, administered twice daily for 10 to 14 days) compared to the protocols used in the other trials.

In the TRICS BASIC trial, remote ischemic conditioning (RIC) was induced surgically via transient clipping of the femoral artery. While this method is commonly employed in preclinical studies ^11^, it significantly differs from the non-invasive approach of external arterial compression used in acute stroke patients ^13^. This fundamental distinction may represent a critical factor contributing to the observed discrepancies in efficacy outcomes between preclinical and clinical studies of this treatment.

Comparison with other studies underscores the complex and context-dependent efficacy of RIC. Unlike the SPAN trial, which terminated the RIC arm for futility ^26^, the TRICS BASIC trial demonstrated a modest yet clearly quantifiable benefit. Notably, the SPAN trial employed an external compression device rather than surgical clipping. Therefore, for the SPAN trial, the same distinction previously highlighted for clinical trials in relation to the TRICS BASIC study remains valid, particularly regarding the method of RIC application, which is non-invasive compared to a surgical approach. Moreover, the SPAN trial focused on long-term outcomes (30 days) using a single behavioral assessment (corner turning test). Conversely, the TRICS BASIC trial emphasized acute outcomes evaluated through a comprehensive neuroscore, potentially offering greater sensitivity for detecting an early treatment effect.

This study has strengths and limitations. Strengths include the rigorous methodology, the large sample size, and the experiments conducted in two species and both sexes. A first limitation is that the primary endpoint was assessed at an acute time point (48 hours), limiting conclusions about the long-term effects of RIC. A second limitation is that we exclusively focused on classical histology of brain samples, while molecular insights into the mechanisms underlying RIC’s neuroprotective effects were not addressed in our study.

This study successfully established a collaborative network among preclinical stroke laboratories, that mirrors clinical trial structures. By refining preclinical trial design, the TRICS BASIC study sets a precedent for future translational research.

In conclusion, the TRICS BASIC trial supports RIC as a promising neuroprotective strategy in acute ischemic stroke and highlights the value of collaborative, rigorously conducted preclinical trials in bridging the gap between experimental research and clinical application.

## Acknowledgments

We thank Davide Carone, Alessandro Versace and Virginia Rogriguez Menendez for their expert advice and technical assistance, and Norberto Oggioni and Luca Crippa for their administrative support. S.B. and C.F. contributed to this work during their personal involvement in the Italian Ministry of University and Research (MUR) “Dipartimenti di Eccellenza 2023-2027”.

## Source of Funding

Italian Ministry of University and Research grant PRIN 2017CY3J3W.

## Disclosures

None.

## Supplemental Material

Figure S1

Figure S2

## Notes

### Competing Interest Statement

The authors have declared no competing interest.

